# LTBP isoforms differentially encode TGF-β spatial localization and activation

**DOI:** 10.64898/2026.08.05.743009

**Authors:** Lixia Wang, Dong Zhou, Jie Tan, Xuezhen Lin, Xinyu Zhao, Pengqing Yuan, Jizhe Liu, Ruxue Li, Niu Wang, Ziying Wang, Fu-Ying Tian, Ying Li, Zhe Zhang, Bo Zhao

**Affiliations:** Zhongshan School of Medicine, Sun Yat-sen University Shenzhen Campus, Sun Yat-Sen University, Shenzhen, Guangdong, 518107, China; State Key Laboratory of Natural and Biomimetic Drugs, Department of Molecular and Cellular Pharmacology, School of Pharmaceutical Sciences, Peking University, Beijing 100191, China; Shenzhen Maternity and Child Healthcare Hospital, Women and Children’s Medical Center, Southern Medical University, Shenzhen, Guangdong, 518028, China; College of Basic Medical Sciences, Zhejiang Chinese Medical University, 310053, Hangzhou, Zhejiang, China; Peking University-Yunnan Baiyao International Medical Research Center, Beijing 100191, China; Center for Life Sciences, Academy for Advanced Interdisciplinary Studies, Peking University, Beijing 100871, China; School of Public Health, Southern Medical University, Guangzhou, Guangdong 510515, China

**Author notes:** These authors contributed equally: Lixia Wang, Dong Zhou, Jie Tan, Xuezhen Lin, Xinyu Zhao.

## Abstract

TGF-β signals through a conserved pathway yet produces diverse outcomes, pointing to regulatory mechanisms upstream of receptor engagement. A family of four latent TGF-β binding proteins (LTBPs) tether pro-TGF-βs and control their localization and activation, but how distinct LTBPs contribute to this regulation is unclear. Here we combine cryo-electron microscopy with functional assays to dissect LTBP-pro-TGF-β interactions. We resolve the LTBP-1/pro-TGF-β1 and LTBP-3/pro-TGF-β3 complex structures, revealing a conserved yet plastic hydrophobic binding interface, and systematically map key residues across all four LTBP and three pro-TGF-β subtypes that determine binding specificity and affinity. Unexpectedly, LTBP-2, previously thought incapable of TGF-β binding, engages pro-TGF-β1 through a covalent linkage via its 16^th^ EGF-like domain, providing a mechanistic link between LTBP-2 function and TGF-β signaling. Beyond tethering, LTBP-3 shifts pro-TGF-β3 from spontaneous activation toward integrin dependence, establishing LTPBs as dynamic modulators that dictate not only where but also how TGF-β is unleashed. Collectively, these findings establish that LTBPs are not redundant ECM scaffolds but a family of functionally distinct regulators that differentially encode TGF-β spatial localization and activation, with implications for isoform-selective therapeutic strategies.

## Introduction

Transforming growth factor beta (TGF-β) is a multifunctional cytokine that plays essential roles in embryonic development, tissue homeostasis, wound healing, and immune regulation^1,2^. Dysregulation of the TGF-β pathway is strongly associated with major human diseases, including cancer, fibrosis, and cardiovascular disorders. Thus, elucidating the mechanisms that regulate TGF-β signaling is critical for understanding disease pathogenesis and developing therapeutic strategies^3–6^.

In mammals, all three TGF-β subtypes (TGF-β1, TGF-β2, and TGF-β3) are synthesized as precursors (pro-TGF-βs)^7^. Pro-TGF-β is comprised of an N-terminal pro-domain (∼250 aa, 37 kDa) and a C-terminal mature growth factor domain (mTGF-β, ∼110 aa, 12.5 kDa)^8^. During biosynthesis, pro-TGF-β covalently associates with specialized “milieu molecules”, including the membrane-bound leucine-rich repeat-containing proteins LRRC32 (GARP)^9^ and LRRC33 (NRROS)^10^, as well as the extracellular matrix proteins known as latent TGF-β binding proteins (LTBP-1, -2, -3, and -4)^11,12^. Milieu molecules determine which microenvironment pro-TGF-β is stored: GARP presents all three pro-TGF-β subtypes on the surface of regulatory T cells (Tregs)^9^, LRRC33 specifically binds pro-TGF-β1 on the surface of myeloid lineage cells^10^, whereas LTBPs localizes pro-TGF-β within the extracellular matrix^12^. Milieu molecules are also essential in the activation of pro-TGF-β by providing mechanical counterforces exerted by integrins ^13,14^. This force-dependent release of mTGF-β from the pro-domain enables its subsequent receptor binding and downstream signaling. Thus, microenvironment-specific localization and activation of pro-TGF-β mediated by milieu molecules represents a key regulatory layer in determining TGF-β’s physiological functions. Targeted blockade of the activation of the disease associated pro-TGF-β-milieu molecule complex is a promising strategy to enhance the specificity and safety of TGF-β therapy.

Previous studies have elucidated the structural basis of how different pro-TGF-β subtypes selectively bind to the membrane-bound leucine-rich repeat-containing proteins) or LRRC33^9,10^. Their distinct binding preferences, and consequently their divergent biological functions, are largely determined by key differences in their prodomain and a specific region of the growth factor domain, particularly the α3 helix^10^. In contrast, despite LTBPs being the first identified class of milieu proteins for TGF-β, the molecular mechanisms by which the three pro-TGF-β subtypes engage the four LTBP isoforms remain poorly understood ^9,10,15^. Unlike the membrane-tethered GARP and LRRC33, LTBPs are secreted ECM proteins that play central roles in ECM organization and ECM-associated pathologies closely related to TGF-β, particularly fibrosis and tumor^12,16,17^.

Among the four LTBP subtypes, LTBP-1 and LTBP-3 have been established as TGF-β binding proteins with distinct functional roles. LTBP-1 is the most extensively studied family member and serves as a key regulator of TGF-β1 bioavailability^18–20^. It covalently associates with the prodomain of all three TGF-β isoforms via its third TGF-β binding (TB) domain, targeting the latent complex to the ECM and providing the mechanical anchor required for integrin-mediated activation^21^. The physiological importance of this LTBP-1/pro-TGF-β1 axis is underscored by *in vivo* studies: mutation of the LTBP-binding cysteine in TGF-β1 pro-domain (C4S) yields mice with inflammatory and tumorigenic phenotypes closely resembling TGF-β1-null animals, demonstrating that covalent association with LTBP is essential for proper TGF-β1 function^17^. The LTBP-1/pro-TGF-β1 axis has been established as a key therapeutic target in renal fibrosis, hepatocellular carcinoma, and other fibrotic diseases^11,12,19,22^. In contrast, LTBP-3 and TGF-β3 are functionally coupled. They converge in specific developmental niches, including developing bone, heart, lung, palate, and skeletal muscle, where they co-orchestrate morphogenetic processes^12,23–26^. This functional coupling is supported by overlapping phenotypes in knockout mice, including cleft palate and skeletal defects^23–25^, and shared genetic links to thoracic aortic aneurysms in humans^27,28^. Beyond development, LTBP-3 is uniquely required among the four LTBP subtypes for adipocyte differentiation, a process specifically driven by TGF-β3 among the three isoforms^29,30^. Further cellular and biochemical evidence includes their interdependent secretion: LTBP-3 is not efficiently secreted by cells on its own^31,32^, but requires co-expression with and binding to TGF-β for secretion, and TGF-β3 secretion from LAG3^+^ Tregs depends on Ltbp3 expression^33–35^.These findings validate selective inhibition of LTBP/pro-TGF-β complexes as a promising therapeutic strategy for fibrosis, cancer, and autoimmune diseases. However, the molecular mechanisms underlying LTBP/pro-TGF-β complex assembly and activation remain poorly understood.

While LTBP–1/pro–TGF–β1 and LTBP–3/pro–TGF–β3 have been established as canonical binding pairs, LTBP-2 presents a striking paradox: although previous studies demonstrated that LTBP-2 does not bind pro-TGF-β through the canonical TB domain, clinical omics data and experimental evidence consistently links LTBP-2 function to TGF-β signaling: elevated LTBP-2 not only serves as a robust prognostic marker for idiopathic pulmonary fibrosis (IPF)^36^, but is also highly enriched in the microenvironment of diverse solid tumors^37^, consistently correlating with TGF-β hyperactivity and poor patient survival^36,38,39^. Depletion of LTBP-2 in cellular and murine models profoundly impairs TGF-β driven disease progression, mitigating both myofibroblast differentiation in fibrosis and pro-tumorigenic processes in cancer^40^. This paradox raises a fundamental question: How does LTBP-2 exert its pro-fibrotic and pro-tumorigenic functions via TGF-β signaling?

Thirdly, LTBPs are conventionally viewed as ECM anchors that not only sequester latent TGF–β in the extracellular matrix but also provide the counterforce required for integrin–mediated mechanical activation. Whether LTBPs play a more active role in controlling pro-TGF–β activation beyond this mechanical function has remained unclear. Among the three pro–TGF–β isoforms, pro–TGF–β1 and pro–TGF–β3 exhibit distinct latency properties: pro–TGF–β1 requires force–dependent activation mediated by integrins^13^, whereas purified pro–TGF–β3 shows a propensity for spontaneous activation in vitro^41^, reflecting its intrinsically weak prodomain latency. This biochemical property raises a physiological puzzle: if pro-TGF–β3 is prone to unregulated activation, how is its activity restrained *in vivo*, given the absence of TGF–β3–driven hyperactivation disorders analogous to those caused by TGF–β1 (such as CED^42^)? Given that LTBP–3 is the primary binding partner of pro-TGF–β3 in the ECM, this raises the question of whether LTBP–3 actively shapes the activation behavior of pro-TGF–β3.

In this study, by combining cryo-electron microscopy (cryo-EM) with comprehensive biochemical, cellular, and functional analyses, we addressed the above fundamental questions. We resolved the cryo-EM structures of LTBP-1/pro-TGF-β1 and LTBP-3/pro-TGF-β3 complexes, defined the structural basis for the covalent and non-covalent interactions underlying their assembly. Through systematic structural, biochemical, cellular and functional analyses, we further delineated the key determinants governing the binding specificity between the three pro-TGF-β subtypes and the four LTBP subtypes. Unlike LTBP-1 and -3, LTBP-2 and -4 lack critical hydrophobic residues in the TB domain, explaining their inability to interact with pro-TGF-β via the canonical mode. Strikingly, we discovered that LTBP-2 covalently binds pro-TGF-β1 through its 16^th^ EGF-like domain. This provides concrete molecular evidence linking LTBP-2’s physiological functions directly to TGF-β signaling, including LTBP-2’s paradoxical role in fibrosis. We further demonstrated that LTBP-3 actively suppresses the spontaneous auto-activation of pro-TGF-β3 by occluding type I receptor access while preserving type II receptor engagement, thereby redirecting it toward integrin-dependent activation.

Collectively, by comparing structures and binding properties across all four LTBPs and three pro–TGF–β subtypes, we uncover a molecular logic that partitions the LTBP family into canonical binders (LTBP-1 and LTBP-3), a non–canonical binder (LTBP-2), and a non–binding member (LTBP-4). Beyond this classification, we show that LTBP engagement actively shapes the activation modality of distinct pro-TGF–β isoforms. These findings establish that LTBPs are not passive ECM scaffolds but rather a family of functionally distinct regulators that collectively orchestrate TGF–β spatial localization and activation specificity. Given the central role of TGF–β dysregulation in fibrosis, cancer, and immune disorders, this functional stratification provides a molecular framework for understanding how distinct LTBP-TGF–β axes contribute to specific pathologies and offers potential structural entry points for isoform–selective therapeutic intervention, an approach that could circumvent the on–target toxicities associated with global TGF-β blockade.

## Results

### 1. Conserved architecture and local plasticity dictate isoform-specific LTBP/pro-TGF-β assembly

To understand how the LTBP family achieves both stable extracellular anchoring and isoform selectivity of pro-TGF-β, we determined the cryo-EM structures of the canonical complexes of LTBP-1/pro-TGF-β1 and LTBP-3/pro-TGF-β3 at resolutions of 3.6 Å and 3.4 Å, respectively (Figures 1A and 1B, S1–S3, and Table S1). These two complexes represent the predominant physiological pairings of the LTBP-TGF-β axis, as supported by extensive biochemical, cellular, and developmental evidence^11,27,33,35,43–45^. Both complexes adopt a 2:1 stoichiometry, with a single LTBP molecule bound to two pro-TGF-β monomers (Figures 1A and 1B, S1-S3). Structural analysis revealed that the LTBP-1/pro-TGF-β1 complex is stabilized through two distinct interactions: a covalent disulfide linkage between pro-domain C4 in each monomer and LTBP-1 C1014/C1039, and a non-covalent interface in which the α3 helix of the mTGF-β1 chain B (mTGF-β1_B_) inserts into a hydrophobic groove on the concave surface of LTBP-1, which is lined with numerous hydrophobic residues (Figures 1D and 1E, S3E-S3G). Hydrogen-deuterium exchange mass spectrometry (HDX-MS) supported the cryo-EM structure by showing that the N-terminal association region and the mTGF-β1_B_ α3 helix exhibit significantly reduced deuterium exchange in the LTBP-1/pro-TGF-β1 complex compared with free pro-TGF-β1(Figure S4C). This result directly maps the solution-state interaction footprint onto our structure and highlights the stabilization of these interfaces upon complex formation.

**Figure 1.**
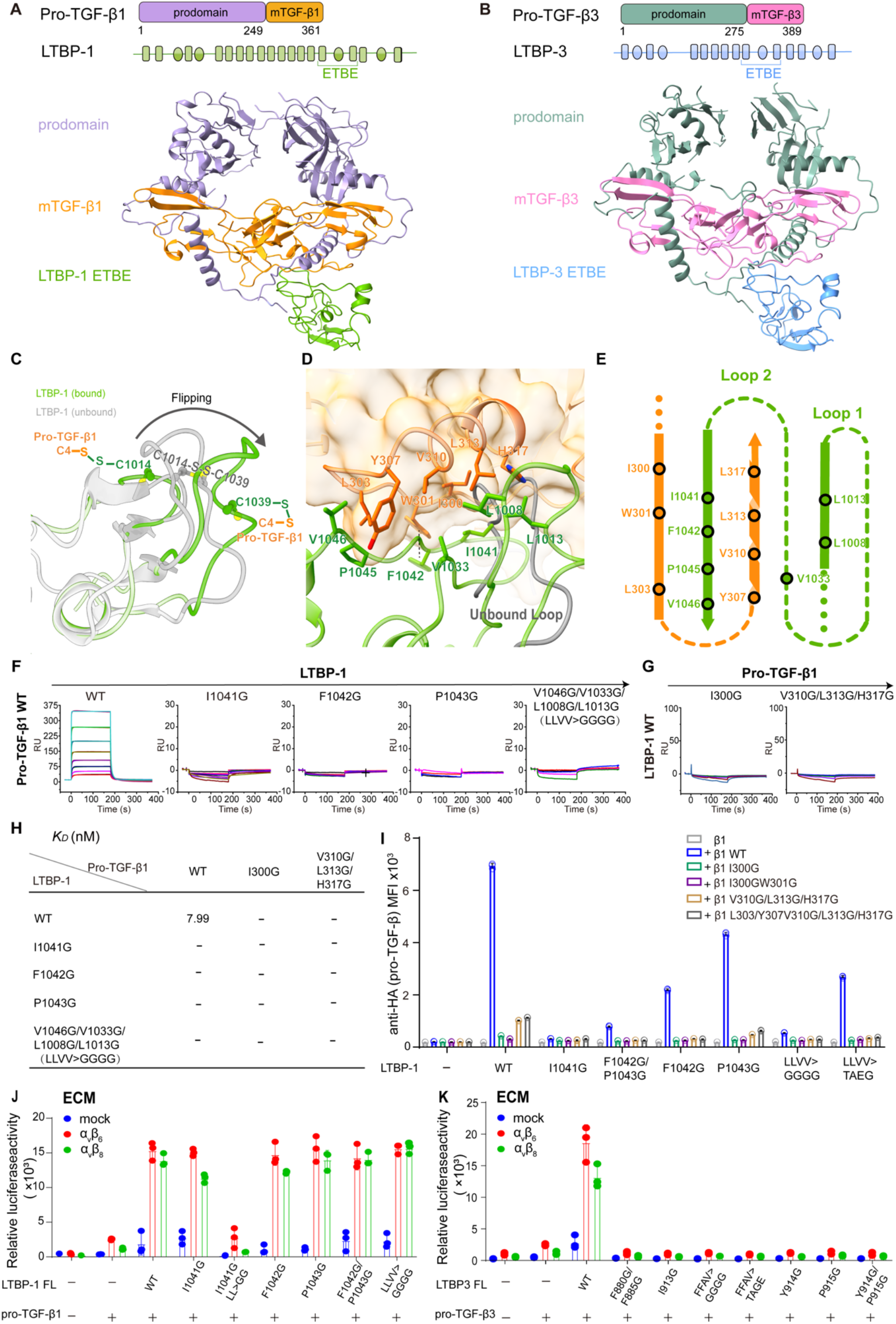
Loop LNDASL Flipping-Mediated Non-Covalent Hydrophobic Interactions Are Crucial for LTBP/pro-TGF-β Complex Formation. (A) Cryo-EM map of the LTBP-1/pro-TGF-β1 complex. (B) Cryo-EM map of the LTBP-3/pro-TGF-β3 complex. (C) Structural comparison of the LNDASL loop in unbound and pro-TGF-β1-bound LTBP-1. (D and E) Hydrophobic interaction interface mediated by loop LNDASL flipping and key hydrophobic residues within the LTBP-1/pro-TGF-β1 complex. (F and G) Surface plasmon resonance (SPR) affinity analysis of pro-TGF-β1 binding to LTBP-1 mutants targeting key hydrophobic residue. (H) Summary of dissociation constants (K_D_) derived from SPR analysis. (I) Flow cytometry quantification of cell-surface pro-TGF-β1 (anti-HA detection) in transfected Expi293F cells, shown as mean fluorescence intensity (MFI). Data represent mean ± s.d. of triplicate measurements from three biologically independent experiments. (J) Effects of LTBP-1 mutants on pro-TGF-β1 activation from ECM. K. Mutations within LTBP-3 diminish its capacity to mediate integrin-dependent activation of pro-TGF-β3 from ECM.

Pro-TGF-β1 maintains a conserved overall architecture when bound to LTBP-1, LTBP-3, GARP (PDB: 6GFF), or LRRC33 (PDB: 7Y1R) (Figure S3C). Despite this global similarity, detailed structural comparisons between the LTBP-1/pro-TGF-β1, pro-TGF-β1/LTBP-3, pro-TGF-β1/GARP, and pro-TGF-β1/LRRC33 complexes reveal significant local plasticity that enables pro-TGF-β1 to adapt to different milieu molecules. First, upon binding to any of these partners, the N-terminal α1 helix of pro-TGF-β1 partially unwinds, while remaining anchored via the conserved disulfide bond (Figure S4A). Second, the α3 helix (residues 307-317) of mTGF-β1_B_ undergoes a ∼48° rotation, accompanied by substantial structural rearrangement of the preceding loop (residues 296-306). These conformational changes enable more optimal packing against the concave surface of LTBP-1 and LTBP-3 (Figures S4B and S4F). Notably, while the α3 helix of mTGF-β1_B_ serves as a common binding interface for all four partners, the specific residues involved differ markedly (Supplementary Figures. 4d-f), resulting in distinct non-covalent interface areas: 972.0 Å² with LTBP-1, 876.1 Å² with LTBP-3, 602.3 Å² with GARP, and 925.6 Å² with LRRC33 (Figures S4D–S4F). Such conformational plasticity allows pro-TGF-β1 to preserve its core architecture while engaging distinct scaffold proteins through tailored local interfaces.

On the LTBP part of the LTBP/pro-TGF-β complex structures, the TB domain and an adjacent EGF-like region of LTBP were clearly resolved (Figures 1A and 1B). Superposition of the bound LTBP-1 structure onto its unbound form (PDB: 1KSQ) shows high overall similarity (Cα RMSD = 1.5 Å), with two significant structural rearrangements (Figure 1C). First, a disulfide-bond switch occurs. In unbound LTBP-1, eight conserved cysteines in the LTBP-1 TB domain form four intrachain disulfide bonds (C1-C3, C2-C6, C4-C7, C5-C8), a pattern conserved across LTBP-2, -3, -4. Upon pro-TGF-β1 binding, cysteines C2 (C1014) and C6 (C1039) instead form interchain disulfides with C4 of each pro-domain monomer (Figures 1C, S3F–S3G). Second, a pronounced conformational change which we named “ LNDASL loop flipping” takes place. In the unbound state, this loop sterically clashes with the α3 helix of mTGF-β1_B_. Upon complex formation, the loop flips outward, shifting the Cα atoms of the central D1010 and A1011 by up to 12 Å. This movement eliminates the clash and creates a hydrophobic pocket lined by LTBP-1 loop1 (L1008-L1013) and loop2 (V1033-V1046) (Figure 1C), into which the mTGF-β1_B_ segment (P296-H317) inserts. Additional stability is conferred by a CH/π-stacking interaction between pro-TGF-β1 W301 and LTBP-1 P1043 (Figures 1D and 1E). Together, these rearrangements provide the critical structural basis for complex assembly. In the LTBP-3/pro-TGF-β3 complex, beyond the covalent disulfide bond linking pro-TGF-β3 (C4) to LTBP-3 (C2/C6), a non-covalent interface (876.1 Å²) was formed. Here, the mTGF-β3_B_ segment (P324-L345) inserts into a hydrophobic pocket delineated by LTBP-3 loop1 (F880-F885) and loop2 (A905-V918) (Figures S8A and S8B). HDX-MS analysis indicated that the mTGF-β3 α3 helix becomes detectable only when bound to LTBP-3, demonstrating that complex formation stabilizes this otherwise flexible region and confirming its role as a key interaction interface (Figure S4C).

To validate the functional importance of the LNDASL loop flipping-mediated hydrophobic interface, we first performed biochemical binding assays at the protein level. Surface plasmon resonance (SPR) measurements showed that mutations in pro-TGF-β1 (I300G, V310G/L313G/H317G) and LTBP-1 (I1041G, F1042G, P1043G, L1008G/L1013G/V1033G/V1046G) severely impaired their binding affinity (Figures 1F–1H). We next developed a GPI-anchored cell-surface display system to evaluate complex formation at the cellular level. In this system, a GPI anchor fused to the C-terminus of the third TB domain of LTBP tethers it to the cell surface (Figures S5A and S5B), allowing flow cytometric detection of pro-TGF-β bound to surface-displayed LTBP (Figure S5C). Disruption of the covalent disulfide linkage between pro-TGF-β3 and LTBP-3 markedly reduced complex formation (68.6%→5.21%) (Figures S5C-S5E), consistent with impaired secretion of cysteine mutants (Figure S5F). The system also reproduced previous findings that D39A and a poly-acidic mutant (E5A/D12A/D17A/D39A/E42A) severely impair complex formation (Figure S6), thereby validating its utility for probing key interfacial contributions. Using this system, we confirmed that hydrophobic residues on both pro-TGF-β1 (I300G, I300G/W301G, V310G/L313G/H317G) and LTBP-1 (I1041G, F1042G, P1043G, L1008G/L1013G/V1033G/V1046G) markedly reduced complex formation, consistent with our SPR results. Combining mutations on both partners completely abolished LTBP-1/pro-TGF-β1 complex assembly (Figures 1I and S7A). In the LTBP-3/pro-TGF-β3 complex, mutational studies identified critical residues in both pro-TGF-β3 (L328G, L328G/R329G, V338G/L341G/L345G) and LTBP-3 (F880G/F885G, F880G/F885G/A905G/V918G, I913G, Y914G/P915G), severely disrupted complex assembly (Figures S8C and S8D). Importantly, all LTBP-GPI mutants exhibited comparable cell surface expression, ruling out expression artifacts as a confounding factor (Figures S7B and S7C).

Finally, we assessed the functional consequences of disrupting this hydrophobic interface on TGF-β activation. Combined mutations in LTBP-1 (I1041G and L1008G/L1013G) reduced pro-TGF-β1 activation, confirming that the LTBP-1/pro-TGF-β1 complex is likewise essential for its activation (Figure 1J). Similarly, mutagenesis of LTBP-3 residues required for complex formation with pro-TGF-β3 (e.g., F880G/F885G, F880G/F885G/A905G/V918G, I913G, Y914G/P915G) severely impaired integrin-dependent activation (Figures 1K), underscoring the indispensable role of LTBP-3. All LTBP mutants showed expression levels comparable to those of full-length LTBP WT, and pro–TGF–β levels were similar across all groups in the activation assay. (Figures S9E and S9F). Specific mutations within pro-TGF-β1 (I300G/W301G causing autoactivation; V310G/L313G/H317G abolishing activity) and pro-TGF-β3 (L328G, L328G/R329G, V338G/L341G/L345G disrupting autoactivation) were found to directly influence intrinsic activity (Figures S9A and S9B). Structural mapping of these residues onto the pro-TGF-β/TβRI (PDB: 3KFD; PDB: 2PJY) complex revealed that I300 and W301 in pro-TGF-β1, together with the corresponding L328 and R329 in pro-TGF-β3, localize to the type I receptor (TβRI) binding interface of the growth factor. This positional overlap indicates that mutations at these sites may directly influence receptor engagement and downstream signaling, in addition to their established effects on partner selectivity. Accordingly, this study focused only on the effects of LTBP-1 and LTBP-3 mutations on activation of pro-TGF-β1 and pro-TGF-β3.

Collectively, these structural, biochemical, cellular, and functional data define a canonical binding strategy shared by LTBP-1 and LTBP-3 via their TB domains, wherein a hydrophobic interface and a covalent disulfide linkage cooperatively sequester pro-TGF-β1 and -β3 while concurrently regulating their integrin-mediated activation.

### 2. Binding Specificity Differences and key Determinants among Three pro-TGF-β Subtypes and Four LTBP Subtypes

To address whether the canonical binding strategy shared by LTBP-1 and LTBP-3 extends to all family members and all pro-TGF-β subtypes, we first performed a comprehensive pairwise interaction screen. Using a GPI-anchored cell-surface display system coupled with western blot analysis, we tested the TB domains of all four LTBPs against each of the three pro-TGF-β isoforms. The results revealed a clear dichotomy: the TB domains of LTBP-1 and LTBP-3 formed covalent complexes with pro-TGF-β1, -β2, and -β3 indiscriminately, whereas LTBP-2 and LTBP-4 failed to engage any of the three pro-TGF-β subtypes under identical conditions (Figures 2C and S10). This discrete binding pattern prompted us to dissect its structural basis by systematically interrogating both sides of the interaction interface.

**Figure 2.**
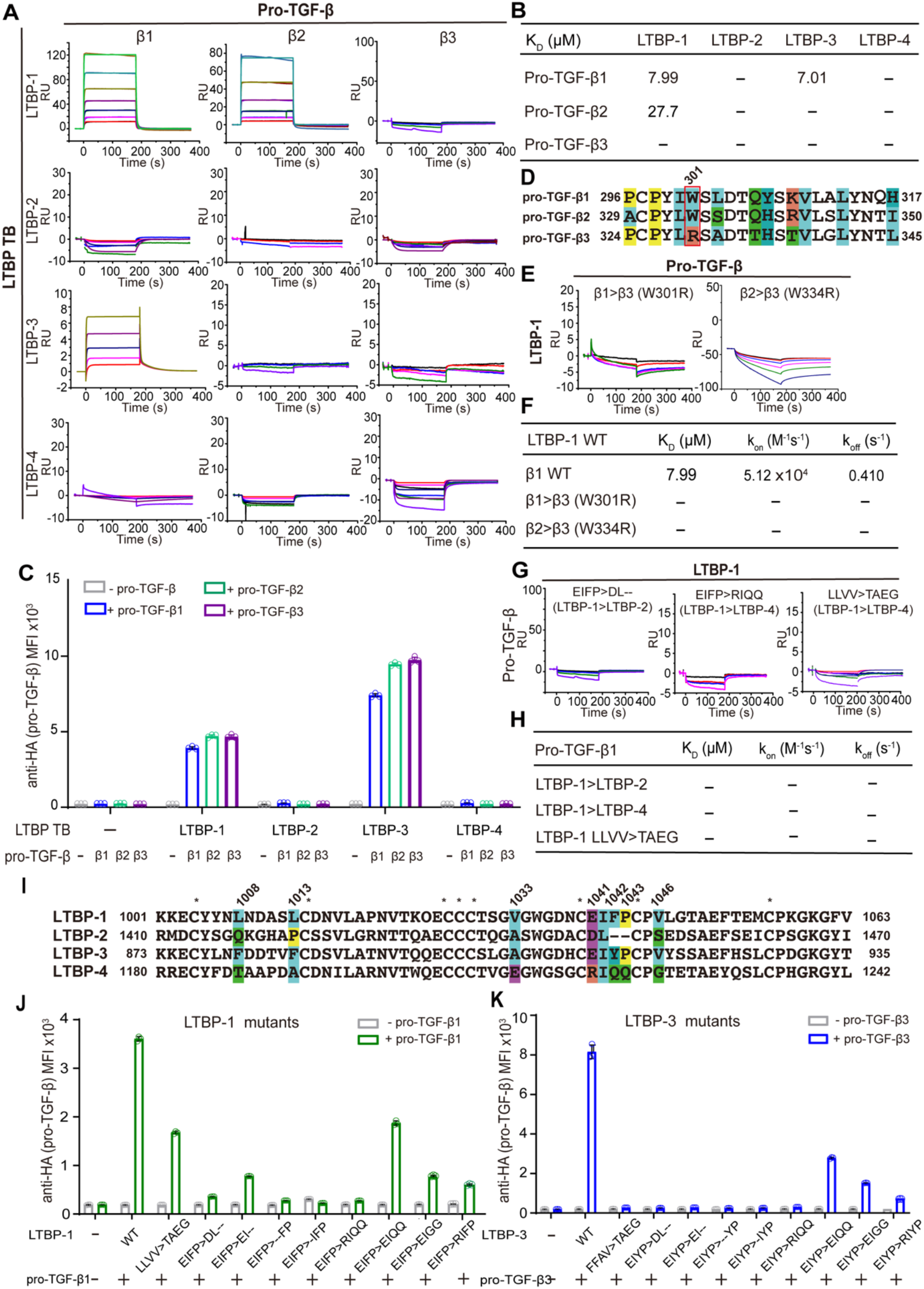
Binding differences of three pro-TGF-β subtypes to four LTBP subtypes. (A and B) Non-covalent binding affinities between pro-TGF-β1, -β2, -β3 and LTBP-1–4. (C) Surface expression of pro-TGF-β1 (detected by anti-HA antibody) on transfected Expi293F cells, measured by flow cytometry. Data shown as mean fluorescence intensity (MFI). (D) Sequence alignment of the α3 helix region in the mTGF-βB of three pro-TGF-β subtypes. (E and F) Contribution of residue W301 to the binding differences between pro-TGF-β3 and -β1, - β2. (G and H) Key residues responsible for the weaker binding affinity of LTBP-2 and LTBP-4 compared to LTBP-1. I. Sequence alignment of the TB3 domain of LTBP-1-4 subtypes. (J and K) MFI of surface pro-TGF-β (detected by anti-HA antibody) on Expi293F cells transfected with the indicated constructs. All experiments were performed in triplicate (n = 3 biologically independent experiments; mean ± s.d.).

To explain why LTBP-2 and LTBP-4 fail to engage pro-TGF-β altogether, we compared the TB domain sequences of the four LTBP family members. Comparative sequence combined with structural analyses revealed that LTBP-1 and LTBP-3 share two classes of conserved motifs within loop1 and loop2 that are entirely absent or altered in LTBP-2 and LTBP-4 (Figure 2I). The first is a previously unrecognized hydrophobic core spanning loop1 and loop2-“LLVV” (L1008/L1013/V1033/V1046) in LTBP-1 and “FFAV” (F880/F885/A905/V918) in LTBP-3-which is positioned at the interaction interface with pro-TGF-β (Figures 1D and S8). The second is a contiguous motif “EIFP” (residues 1040-1043) in LTBP-1 and“EIYP” in LTBP-3, located within loop2, which has been previously reported^46^, but whose mechanistic basis whose structural basis for pro–TGF–β engagement has remained unclear.

To verify the importance of individual regions for pro–TGF–β binding. We first mutated the hydrophobic core in LTBP-1 (LLVV>TAEG) or LTBP-3 (FFAV>TAEG) to match the LTBP-4 sequence, and found that it completely abolished complex formation (Figures 2G-2K and S11). Similarly, replacing the ”EIFP” motif in LTBP-1 with corresponding sequences from non-binding LTBP-2 (EIFP>DL--) or LTBP-4 (EIFP>RIQQ) eliminated binding affinity (Figures 2G and 2H). We further dissected the contributions of individual structural elements within the ”EIFP” motif. Systematic truncations (EIFP>EI/IFP/FP) and hydrophobicity-disrupting mutations (EIFP>EIQQ/EIGS) progressively impaired complex formation, underscoring the critical importance of both motif length and the terminal hydrophobic ”FP” residues (Figures 2J). Parallel mutations in LTBP-3’s “EIYP” motif (EIYP>DL--, EIYP>RIQQ, EIYP>EI, EIYP>IFP, EIYP>FP, EIYP>EIQQ, EIYP>EIGG) produced consistent defects (Figures 2K and S11). Furthermore, a single charge-reversal mutation substituting glutamate with arginine (EIFP>RIFP in LTBP-1; EIYP>RIYP in LTBP-3) to mimic the LTBP-4 sequence markedly reduced complex formation (Figures 2J-2K and S11), highlighting the importance of the specific electrostatic complementarity interaction that is uniquely present in LTBP-1 and -3 but absent in LTBP-4. Together, these results demonstrate that the binding competence of LTBPs is determined by the presence of both conserved motifs: the hydrophobic core (“LLVV/FFAV”) and the full-length ”EIFP/EIYP” motif. The absence of critical structural elements in these motifs in LTBP-2 and LTBP-4 directly accounts for their failure to bind any pro-TGF-β isoform.

The LTBP motifs essential for binding are functionally coupled to pro–TGF–β activation. To determine whether these motifs carry functional relevance beyond complex assembly, we assessed their impact on integrin–mediated pro–TGF–β activation. Individual glycine substitutions within the LLVV and EIFP motifs of LTBP-1 (L1008G/L1013G, I1041G) and the FFAV and EIYP motifs of LTBP-3 (F880G/F885G, I913G, Y914G/P915G) not only disrupted complex assembly but also impaired integrin-dependent activation (Figures 1J and 1K). This establishes that the same hydrophobic interface residues governing binding competence are also essential for the activation function of LTBP-1 and LTBP-3.

Having established the functional importance of the TB–domain–mediated interface for LTBP–1 and LTBP–3, we next asked whether LTBP–2 and LTBP–4, which lack the conserved binding motifs, would similarly fail to support activation (Figure S12). We compared full length LTBP-1, -2, -3, and -4 for their ability to anchor and activate pro-TGF-β in the ECM. As expected, LTBP-4 failed to support ECM localization and activation of pro-TGF-β, consistent with its complete inability to engage its binding partner through the TB domain. Unexpectedly, despite lacking detectable TB-domain-mediated binding, LTBP-2 retained the capacity to anchor pro-TGF-β in the ECM and activated it. This dissociation between TB-domain binding and functional activity strongly suggests that LTBP-2 may promote pro-TGF-β activation through a mechanism independent of its TB domain, potentially involving other regions of the protein or indirect interactions (Result section 3).

Having established the LTBP-side determinants that govern the binary “bind or not bind” decision, we next asked whether the three pro-TGF-β isoforms exhibit any binding preferences among themselves, a question that could only be addressed after confirming that LTBP-1 and -3 are capable of binding all three. Although the cysteines responsible for disulfide-linked covalent conjugation are strictly conserved across all pro-TGF-β isoforms, we reasoned that non-covalent interactions might confer subtle differences in binding efficiency. To test this, we measured the non-covalent affinity of each pro-TGF-β subtype for LTBP-1 using surface plasmon resonance (SPR) (Figures 2A and 2B). While pro-TGF-β1 and -β2 bound LTBP-1 with comparable dissociation constants (K_D_ ∼10⁻⁵ M), pro-TGF-β3 exhibited a markedly weaker affinity that fell below the detection limit of the assay. Sequence alignment at the LTBP/pro-TGF-β binding interface pinpointed a single non-conserved residue: arginine at position 329 in TGF-β3, which is replaced by a conserved tryptophan in both TGF-β1 and TGF-β2 (Figure 2D). Mutagenesis confirmed this residue as a critical specificity switch. Introducing the β3-specific arginine into β1 (W301R) or β2 (W334R) severely impaired their non-covalent affinity for LTBP-1 (Figures 2D-2F). Conversely, replacing the β3-specific R329 with the β1/β2-conserved tryptophan (R329W) was not detectable by SPR, likely due to the technique’s inherent limitations in resolving weak interactions (K_D_ > 10⁻⁴∼10⁻⁵ M). Collectively, these results identify W301 (conserved in β1/β2) and R329 (β3-specific) as key determinants of differential non-covalent binding efficiency between pro-TGF-β3 and its β1/β2 counterparts. Although non-covalent interactions of TGF-β3 to LTBP was weaker. cell-surface display data unambiguously showed that LTBP-3/pro-TGF-β3 complex formation was as robust as the LTBP-1/pro-TGF-β1 pair (Figures S10A, S10C and S10E). Suggesting the weakened non-covalent interaction of pro-TGF-β3 did not affect its covalent disulfide linkage to LTBPs, suggesting a functional division of labor: the non-covalent interface primarily governs the efficiency and kinetics of initial complex assembly, whereas the covalent bond ensures stable sequestration of the latent complex in the extracellular matrix. Thus, in contrast to the LTBP-side determinants that dictate binary binding competence, the pro-TGF-β-side determinants modulate the efficiency of complex formation without affecting the final covalent outcome.

In summary, our systematic dissection establishes a two-layered model of binding specificity. On the LTBP side, the presence of conserved hydrophobic and charged motifs (“LLVV/FFAV” and “EIFP/EIYP”) determines binary binding competence: LTBP-1 and -3 possess them and therefore bind all pro-TGF-βs, whereas LTBP-2 and -4 lack them and therefore bind none. On the pro-TGF-β side, a single residue difference (W301 in β1/β2 versus R329 in β3) modulates binding efficiency, affecting non-covalent affinity and assembly kinetics without compromising the final covalent complex. These findings reveal that isoform-specific binding between LTBPs and pro-TGF-βs is governed by qualitatively distinct determinants on each interacting partner, with functional consequences that extend to TGF-β activation, and point to an alternative activation mechanism for LTBP-2 that warrants further investigation.

### 3. LTBP-2 Binds and Activates Pro-TGF-β1 via its 16^th^ EGF-like Domain

The finding that LTBP–2, despite lacking TB–domain–mediated binding, retained the capacity to activate pro–TGF–β in our full–length activation assays suggests that LTBP–2–mediated activation potentially operates through a mechanism distinct from that of LTBP–1 and LTBP–3, one that does not strictly require TB–domain–dependent binding.

To investigate the alternative mechanism by which LTBP-2 engages pro-TGF-β1, we generated a series of truncation mutants and assessed their binding capacities. Cell-surface binding assays and pull-down analyses consistently demonstrated that, among the four truncations tested (LTBP-2 N-terminal comprising of 1^st^-3^rd^ EGF-like, and TB2 domains, 4^th^-15^th^ EGF-like; ETBE comprising of 16^th^ EGF-like, TB3 and 17^th^ EGF-like domains; and C-terminal comprising of 18^th^-20^th^ EGF-like and TB4 domains), only the ETBE fragment formed a prominent high-molecular-weight covalent complex with pro-TGF-β1, whereas the other three truncations showed no detectable binding (Figure S13). These results indicate that the ETBE region is both necessary and sufficient for LTBP-2 to engage pro-TGF-β1 through a covalent mechanism. We further subdivided the ETBE fragment into its constituent domains: the 16^th^ EGF–like, TB, and TBE (TB3 and 17^th^ EGF-like domains) domains. Through complementary truncation and pull–down analyses, we identified that the isolated 16^th^ EGF–like domain of LTBP–2 is solely sufficient to form a distinct covalent complex with pro–TGF–β1, whereas the adjacent TB and TBE domains completely failed to bind pro–TGF–β1 (Figures 3C), definitively mapping the interaction site to the 16^th^ EGF–like module. Furthermore, this covalent capture mechanism is highly unique to LTBP–2, as the corresponding EGF–like domains from LTBP–1, –3, or –4 showed no significant complex formation with pro–TGF–β1 (Figures S14D-14F).

**Figure 3.**
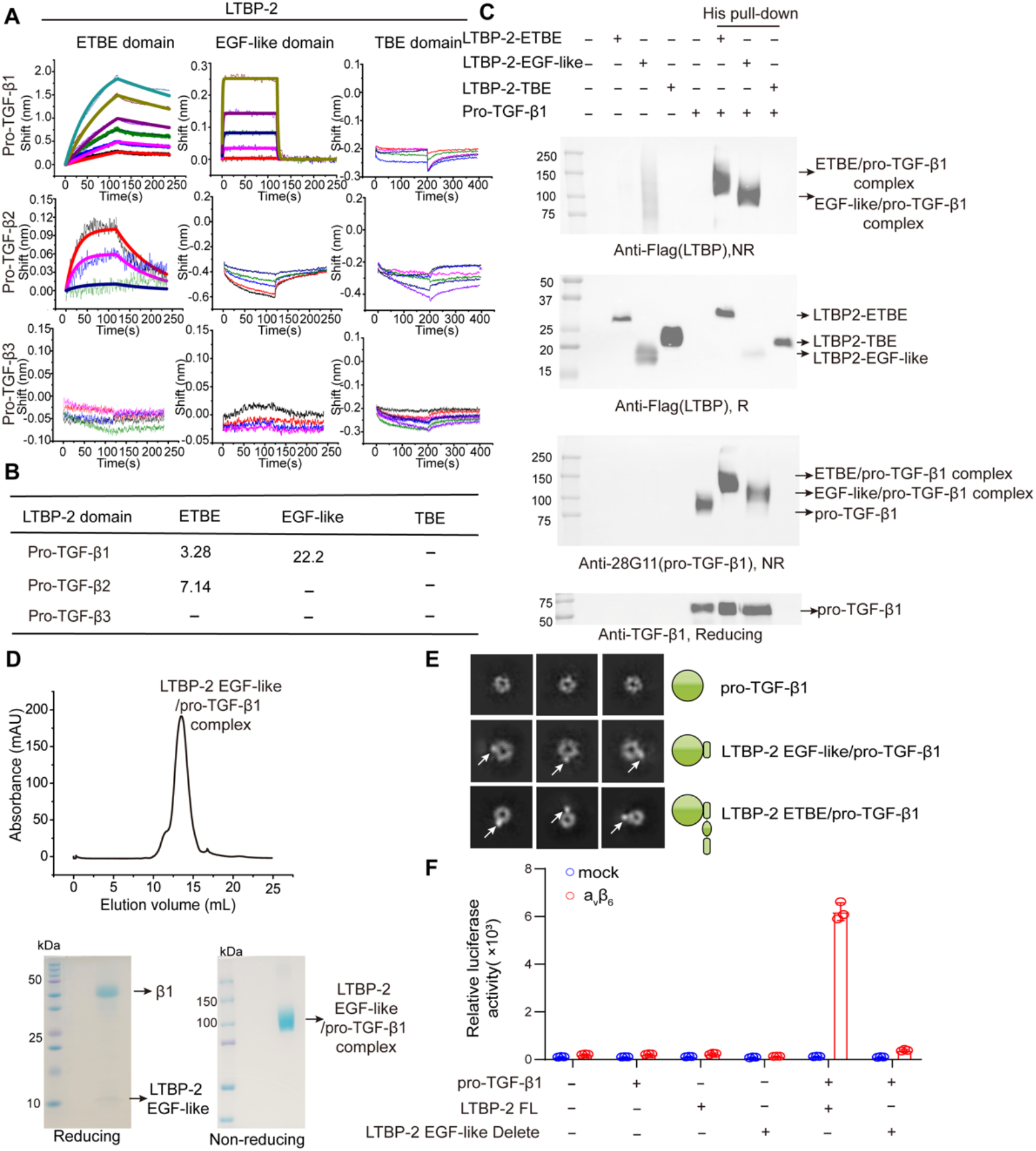
The 16^th^ EGF-like domain of LTBP-2 mediates covalent binding and activation of pro-TGF-β1. (A) Binding affinities of different LTBP-2 domains for pro-TGF-β subtypes (β1, β2, β3). (B) Summary of dissociation constants (KD) and kinetic parameters (kon, koff) corresponding to the interactions shown in (A). (C) Pull-down analysis of LTBP-2 domain interactions with pro-TGF-β1. (D) Size-exclusion chromatography (SEC) purification of the complex formed between pro-TGF-β1 and the 16^th^ EGF-like domain of LTBP-2. (E) Representative negative-stain electron microscopy class averages of pro-TGF-β1 alone and in complex with the LTBP-2 16^th^ EGF-like or ETBE domains. (F) Deletion of the 16^th^ EGF-like domain impairs LTBP-2-mediated pro-TGF-β1 activation.

Having established the 16^th^ EGF–like domain as the covalent anchor, we next evaluated its isoform selectivity. Cell–surface display assays revealed that the LTBP–2 16^th^ EGF–like domain binds robustly to pro–TGF–β1, but only weakly interacts with s pro–TGF–β2 and –β3 (Figures S15A-14C). Consistent with this cell–based finding, rigorous non–covalent affinity measurements by BLI demonstrated that the LTBP–2 16^th^ EGF–like domain binds to pro–TGF–β1 with an apparent K_D_ of 22.2 μM, while exhibiting no detectable binding to pro–TGF–β2 or –β3 (Figures 3A and 3B). Collectively, these results indicate that LTBP–2 binding to pro–TGF–β1 is mediated predominantly via covalent interactions involving its 16^th^ EGF–like domain, a strategy fundamentally distinct from the TB–domain–dependent mechanism utilized by LTBP–1 and LTBP–3.

To investigate the molecular basis of LTBP-2-mediated pro-TGF-β1 activation, we purified complexes of the LTBP-2 16^th^ EGF-like domain with pro-TGF-β1 (Figure 3D). Both complexes eluted as stable, monodisperse peaks during gel filtration and were subsequently analyzed by cryo-EM. 2D classification revealed that pro-TGF-β1 adopts its characteristic ring-like architecture in both complexes, with an additional punctate density at the periphery-likely corresponding to the bound LTBP-2 fragments (Figures 3E, S16). Despite extensive efforts, we were unable to obtain a high-resolution 3D reconstruction of either complex. This most probably arose from the inherent flexibility of the LTBP-2 fragments relative to the pro-TGF-β1 ring, which generates continuous conformational heterogeneity and prevents consistent particle alignment. As a result, the peripheral density attributed to LTBP-2 was observed only in some 2D class averages and became progressively less well defined during 3D classification and refinement. In the final refined map, only a residual extra-structural density extending from pro-TGF-β1 was discernible (dashed red circles, Figure S16), nevertheless confirming the presence of a EGF-like fragment despite the lack of a well-resolved structure.

Functional assessment showed that deletion of the 16^th^ EGF-like domain significantly impaired activation of pro-TGF-β1 by LTBP-2 (Figures 3F, S12E and S12F). These biochemical, structural and functional data collectively indicate that domain-specific binding is a key mechanism by which LTBP-2 modulates the physiological and pathological functions of pro-TGF-β1 *in vivo*, expanding the functional repertoire of the LTBP family beyond the canonical TB-domain binding mode.

### 4. LTBP-3 Reprograms pro-TGF-β3 Activation from Autoactivation to Integrin-Dependence

It has been previously reported that, unlike pro–TGF–β1 and -β2, which are intrinsically latent and require integrins and milieu molecules such as LTBPs for activation, pro–TGF–β3 exhibits intrinsically weak latency and a propensity for spontaneous auto–activation. This raises the question of whether LTBP–3 merely functions as a spatial sequester for pro–TGF–β3. Surprisingly, in our activation assays, we observed that when pro–TGF–β3 is covalently associated with LTBP–3 and anchored in the ECM, its activation is totally integrin–dependent activation (Figures 4F-4H). This finding prompted us to investigate how LTBP–3 reshapes the activation modality of pro–TGF–β3 and whether LTBP–3 serves as a critical switch that overrides the intrinsic auto–activation propensity of β3, coupling its activation to integrin–mediated mechanical signaling.

To address this, we first assessed the intrinsic latency of various TGF-β forms using a (CAGA)₁₂-luciferase reporter system with serial dilutions of purified proteins (Figure 4A). The panel comprised mature TGF-β isoforms (mTGF-β1, -β2, and -β3), full-length pro-TGF-β isoforms (pro-TGF-β1, -β2, and -β3) with intact furin cleavage sites, a furin-site mutant of pro-TGF-β3 (pro-TGF-β3-FM), and LTBP-3/pro-TGF-β3 complex. Compared with pro-TGF-β1 (EC₅₀ > 100 nM) and pro-TGF-β2 (EC₅₀ = 0.12 nM), pro-TGF-β3 exhibited a distinct propensity for auto-activation (EC₅₀ = 0.03 nM), approaching the potency of mature mTGF-β3 (EC₅₀ = 0.01 nM). This observation underscores a subtype-specific difference in latent complex regulation. The auto-activation of pro-TGF-β3 was completely abolished by furin-site mutation (EC₅₀ > 100 nM), confirming the essential role of pro-domain dissociation. Notably, the LTBP-3/pro-TGF-β3 complex showed a substantial 177-fold weaker signaling capacity (EC₅₀ = 1.77 nM) relative to free pro-TGF-β3 (Figures 4A), confirming that LTBP-3 binding abolishes auto-activation of pro-TGF-β3.

**Figure 4.**
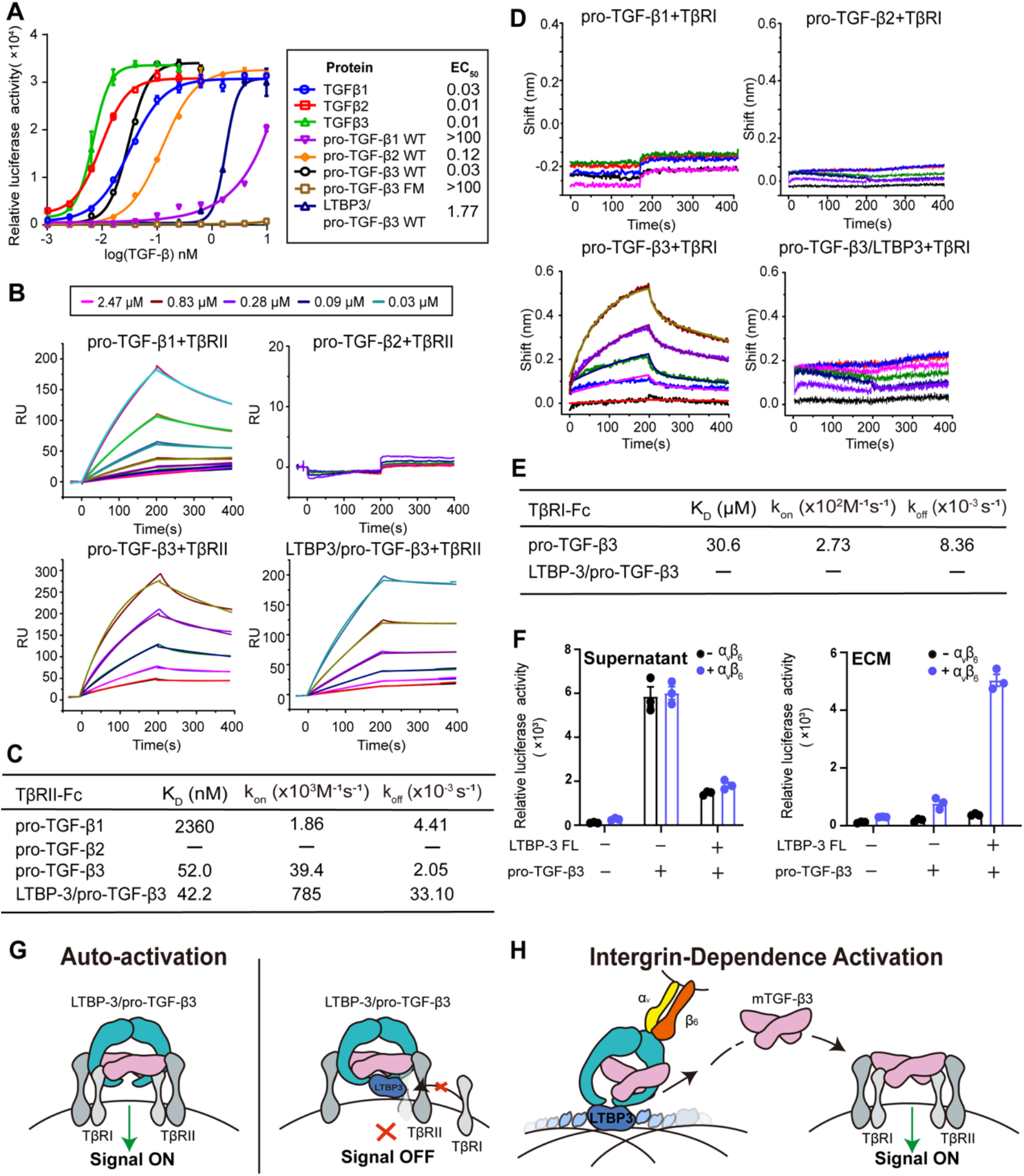
Role of LTBP in pro-TGF-β activation. (A) TGF-β activity measured in (CAGA)_12_-luciferase reporter cells incubated with a series of concentrations of recombinant human pro-TGF-β subtype proteins. (B and C) Surface plasmon resonance (SPR) analysis of binding affinities between pro-TGF-β1, pro-TGF-β2, pro-TGF-β3, the LTBP-3/pro-TGF-β3 complex, and TβRII. (B) Representative sensorgram. (C) Summary of dissociation constants (KD) and kinetic parameters (kon and koff). (D and E) SPR analysis of binding affinity between pro-TGF-β3, or the LTBP-3/pro-TGF-β3 complex and TβRI (D). Summary of dissociation constants and kinetic parameters (E). (F and G) LTBP-3 shifts pro-TGF-β3 activation from autoactivation to an integrin-dependent pathway.

Differential receptor binding underlies the activation mode shift. We systematically compared the binding affinities of purified pro–TGF–β1, –β2, and –β3, as well as the LTBP–3/pro–TGF–β3 complex, to the signaling receptors TβRI and TβRII. We systematically compared the binding affinities of purified pro–TGF–β1, –β2, and –β3, as well as the LTBP–3/pro–TGF–β3 complex, to the signaling receptors TβRI and TβRII. Pro–TGF–β3 exhibited ∼45–fold higher affinity for TβRII (K_D_ = 52 nM) than pro–TGF–β1 (K_D_ = 2360 nM), consistent with their divergent EC₅₀ values, whereas pro–TGF–β2 showed no detectable interaction with TβRII, implying reliance on TβRIII (betaglycan) for signaling (Figures 4B and 4C). Only pro–TGF–β3 could bind TβRI, confirming that its prodomain confers weak latency and don’t block receptor binding of its growth-factor domain. Notably, complex formation with LTBP-3 did not alter the TβRII affinity of pro-TGF-β3 (42.2 nM vs. 52 nM; Fig. 4c d), but completely blocked out TβRI binding (Figures 4D and 4E). Structural analysis revealed that LTBP-3 binding sterically occludes the TβRI docking site (Figures S17A).

Consistent with the biochemical evidence, functional studies showed that when anchored to the ECM or cell surface by LTBP–3, pro–TGF–β3 activation became integrin–dependent (Figures 4F and 4H). Stretching dynamics simulations further supported that LTBP–3 can endure mechanical forces exerted by integrins to induce conformational changes that activate pro–TGF–β3 (Figures S17B and S17C). Together, these data demonstrate that LTBP–3 suppresses spontaneous auto–activation of pro–TGF–β3 by occluding TβRI access, while reprogramming it to integrin–mediated mechanical activation.

This finding establishes LTBP as a decisive factor ensuring that TGF-β3 signaling is not constitutively active *in vivo* but rather restricted to specific microenvironments where integrin-mediated force provides the necessary trigger.

## Discussion

The functional diversity of TGF-β despite its conserved receptor signaling has long pointed to regulation upstream of receptor engagement^5,8^. Extracellular milieu molecules, including membrane-bound GARP/LRRC33 and ECM-resident LTBPs, govern pro-TGF-β localization and activation^9,10,12^. While the structural basis for GARP/LRRC33-mediated pro-TGF-β presentation has been established^9,10^, how LTBPs engage and regulate distinct pro-TGF-β isoforms has remained unclear. Our work addresses this gap by systematically characterizing LTBP-pro-TGF-β interactions across all four LTBP and three pro-TGF-β subtypes, revealing unexpected mechanisms that fundamentally revise the view of LTBPs as passive ECM scaffolds.

Although GARP/LRRC33 and LTBPs adopt markedly different architectures, they converge on a shared binding interface on pro-TGF-β, centered on the α3 helix of the mature growth factor domain and the N-terminal α1 helix of the pro-domain. This interface is intrinsically flexible in the unbound state, as seen in unbound pro-TGF-β1 and integrin-bound structures, and becomes stabilized upon partner engagement, as observed in complex structures with LTBP-1, GARP, and LRRC33. The α3 helix is uniquely exposed beyond the prodomain, making it accessible to diverse milieu molecules. The mechanisms of engagement, however, differ fundamentally. In GARP and LRRC33, pro-TGF-β is presented on the surface of the horseshoe-shaped LRR domain. In LTBPs, engagement requires LNDASL loop flipping, a conformational rearrangement that relieves steric hindrance and enables hydrophobic interactions between LTBP and the α3 helix. This mechanistic distinction explains how a globular TB domain can engage the same α3 helix recognized by an LRR domain, and it highlights a broader principle: other TGF-β family cytokines, such as BMPs^7^, are also synthesized as precursors with exposed α3-like helices and N-terminal free cysteines, suggesting that this mode of extracellular regulation may be evolutionarily conserved.

Beyond serving as a shared docking site, the α3 helix also functions as a selectivity hub that discriminates between partner classes and TGF-β isoforms. Residue-specific determinants at the N-terminal entrance of the helix, such as W301 (critical for LRRC33 binding) and I300 (dominant for LTBP-1 binding), provide structural readouts for distinguishing LRR-domain proteins from TB-domain proteins. In addition, sequence variations within the same interface modulate isoform-specific binding behaviors. The substitution of the conserved tryptophan in pro-TGF-β1/β2 (W301/W334) with arginine in pro-TGF-β3 (R329) weakens non-covalent affinity for LTBPs, yet this does not prevent covalent complex formation. This points to a sequential assembly model in which the α3 helix-mediated non-covalent interactions facilitate initial docking, while the subsequent disulfide linkage ensures stable ECM sequestration. Thus, the α3 helix serves as a central hub that enables both promiscuous engagement and selective encoding, resolving the apparent paradox of how a shared ligand achieves divergent regulatory outcomes.

The LTBP-1/pro-TGF-β1 axis has been extensively characterized as a central regulator of TGF-β bioavailability^13^, with well-established roles in ECM assembly^18^, fibrosis^11,20^, and cancer^2^. In contrast, the functional significance of the LTBP-3/pro-TGF-β3 axis is less understood but is underscored by compelling genetic evidence. LTBP-3 knockout mice develop cleft palate, skeletal defects, and lung malformations, closely phenocopying TGF-β3-null animals^23–25^. Mutations in human LTBP3 cause thoracic aortic aneurysms and dissections, a vascular phenotype indistinguishable from that driven by TGF-β3 mutations^27,28^. Beyond development, LTBP-3 is uniquely required among the four LTBP subtypes for adipocyte differentiation, a process specifically driven by TGF-β3 among the three isoforms^29,30^. These observations point to a dedicated functional link between LTBP-3 and TGF-β3.

Our structural and biochemical analyses, however, reveal that LTBP-3 does not discriminate between pro-TGF-β isoforms at the binding level. Thus, the specificity of the LTBP-3/pro-TGF-β3 axis cannot be explained by intrinsic binding selectivity. Instead, it likely arises from their overlapping spatial expression. LTBP-3 and TGF-β3 converge in specific developmental niches, including bone, heart, lung, palate, and skeletal muscle, where they jointly orchestrate morphogenesis. This co-expression is functionally reinforced by their interdependent secretion: LTBP-3 requires TGF-β binding for its own secretion^31,32^, and TGF-β3 secretion from LAG3^+^ Tregs depends on Ltbp3 expression^33–35^. In the growth plate, LTBP-3 is expressed by chondrocytes at multiple differentiation stages^24,28^. In contrast, LTBP-1 and TGF-β1 co-localize extensively in matrix-remodeling contexts such as dermal elastic microfibrils^15,47^. These distinct spatial logics define the physiological specificity of each LTBP-TGF-β pair.

Beyond co-expression, LTBP-3 adds a regulatory layer that actively shapes TGF-β3 activation. We find that LTBP-3 engagement does not interfere with type II receptor binding but completely occludes type I receptor access. This steric blockade is especially critical for TGF-β3, whose prodomain exhibits weak latency and a propensity for spontaneous auto-activation. By blocking type I receptor engagement, LTBP-3 suppresses unregulated release while preserving the potential for integrin-dependent activation. Thus, the LTBP-3/pro-TGF-β3 axis is defined by two complementary layers of specificity: spatial specificity conferred by co-expression, and activation specificity enforced by receptor occlusion. Together, they ensure that TGF-β3 is delivered to the right place and unleashed only through the right trigger.

Beyond the LTBP-3/pro-TGF-β3 axis, our data reveal a general principle of LTBP-pro-TGF-β interactions: non-covalent affinity does not predict covalent complex formation. The weaker non-covalent interaction of pro-TGF-β3 with LTBPs, despite unaffected covalent attachment, indicates that these two binding modes serve distinct functions. Covalent engagement ensures stable ECM sequestration, whereas the non-covalent interface governs the efficiency and dynamics of initial complex assembly. This division of labor is consistent with the observation that residues required for covalent engagement are conserved between TGF–β3 and β1/β2, whereas residues governing non–covalent affinity, such as W301 and R329, are divergent. This suggests that TGF–β3, unlike β1 and β2, may be subject to distinct assembly dynamics, a possibility that warrants further investigation.

Another major finding of our study is the identification of a non-canonical binding mechanism for LTBP-2. We discovered that LTBP-2 directly binds pro-TGF-β1 through a covalent linkage involving its 16^th^ EGF-like domain. This finding fundamentally revises the prevailing paradigm that LTBP-2 lacks TGF-β binding capacity and resolves a long-standing paradox: although previously assumed incapable of interacting with TGF-β, LTBP-2 nevertheless plays indispensable roles in TGF-β regulation, particularly in idiopathic pulmonary fibrosis (IPF)^36,38^. Unlike other LTBP subtypes that engage latent TGF-β through TB-domain-dependent interactions^21^, LTBP-2 employs a non-canonical binding mode, unveiling a previously unrecognized mechanism of latent TGF-β regulation.

The pathophysiological relevance of this LTBP-2/pro-TGF-β1 axis is strongly supported by functional evidence. In IPF, plasma LTBP-2 levels are markedly elevated in patients with acute exacerbations and correlate with disease severity^38^. Proteomic analyses reveal that LTBP-2 abundance in the ECM deposited by TGF-β1-stimulated IPF fibroblasts strongly correlates with TGF-β activity^39^, and both protein and transcript levels are robustly upregulated in IPF-derived myofibroblasts upon TGF-β1 stimulation^36^. LTBP-2 knockout mice exhibit attenuated TGF-β signaling in experimental fibrosis models^40^. Beyond pulmonary fibrosis, silencing LTBP-2 in human fibroblasts abrogates TGF-β1-induced upregulation of α-SMA, fibronectin, and type I collagen^48^, while antisense-mediated knockdown in rat astrocytes reduces both LTBP-2 expression and TGF-β activity^49^. Collectively, these findings indicate that LTBP-2 functions as a broad-spectrum regulator of latent TGF-β storage and activation within the ECM.

This functional role is further corroborated by human genetic evidence. A naturally occurring mutation in the 16^th^ EGF-like domain of LTBP-2 (C1372R) disrupts disulfide bond formation, compromises structural integrity, and weakens microfibril anchoring^50^. Another pathogenic mutation, Asp1345Glyfs*6^51^, further implicates this domain in disease. Moreover, amino acids 1816-1821 at the LTBP-2 carboxyl terminus, reported as the principal fibrillin-1 binding site, mediate ECM-associated pro-TGF-β regulation; the mutant H1816PfsX28 abolishes this interaction, impairing TGF-β activity and leading to pathogenic consequences^52^. Variant data from multiple databases (LOVD, cBioPortal, ClinVar, and gnomAD) further emphasize that mutations in the 16^th^ EGF-like domain are recurrent and likely deleterious.

It is worth noting that atomic modeling of the LTBP-2 16^th^ EGF-like domain was precluded in our cryo-EM reconstruction by its small size, peripheral location, and conformational flexibility, which produced smeared density insufficient for backbone tracing. Despite this limitation, the combined functional and genetic evidence firmly establishes the 16^th^ EGF-like domain as a critical structural and functional module governing pro-TGF-β1 binding and activation. These findings implicate the LTBP-2/pro-TGF-β1 axis in fibrotic pathology and position LTBP-2 as a potential therapeutic target for fibrotic diseases.

Taken together, our findings establish that the four LTBPs constitute a non-redundant regulatory network in which structural diversification of a shared evolutionary template has produced distinct binding strategies, subtype selectivities, and activation modes. LTBP–1 and LTBP–3 employ a conserved TB–domain mechanism, with differences in pro–TGF–β preferences shaped by expression patterns. LTBP–3 further adds a unique regulatory layer through type I receptor occlusion, suppressing spontaneous auto–activation of TGF–β3 while enabling integrin–dependent activation. LTBP–2 has evolved an alternative covalent mechanism via its 16^th^ EGF–like domain, linking it specifically to pro–TGF–β1–driven fibrotic disease. LTBP–4, in contrast, appears to have lost the hydrophobic interface required for stable non–covalent engagement, potentially dedicating its structural modules primarily to ECM architecture. Thus, the LTBP family has diversified along two axes: LTBP–1, –2, and –3 have evolved distinct strategies for TGF–β regulation, while LTBP–4 may have repurposed its conserved framework toward structural roles. This functional stratification carries direct therapeutic implications: targeting specific LTBP–pro–TGF–β interfaces could disrupt disease–relevant axes while sparing homeostatic functions, with the loop flipping–mediated hydrophobic interface and the 16^th^ EGF–like domain representing candidate targets for isoform–selective intervention.

## Methods

### Protein expression and purification

For pro-TGF-β and LTBP complex, the DNA sequences encoding the full-length pro-TGF-β1 (residues 1-361 excluding signal peptide) and pro-TGF-β3 (residues 1-389 excluding signal peptide) were cloned into the pcDNA3.4 expression vector. The LTBP-1 ETBE truncation (residues 940-1121) and LTBP-3 ETBE domain (residues 823-1031) were cloned into the pd2529 expression vector (ATUM). A 10 × His-tag was attached to the C-terminus of LTBP constructs. For TβRII, the DNA sequence encoding its extracellular domain (residues 1-166 excluding signal peptide) was cloned into the same pd2529 vector with a C-terminal human IgG1 Fc tag.

All plasmids were transiently transfected into Expi293F cells (Thermo Fisher Scientific), which were cultured in OPM-293 CD05 medium (OPM, 81075-001) in suspension, using PEI (Polyethylenimine, linear, MW 25,000) as the transfection reagent at a DNA:PEI ratio of 1:3 (w/w). For the expression of LTBP/pro-TGF-β complex, equal amounts of pcDNA3.4-pro-TGF-β (pro-TGF-β1 or pro-TGF-β3) and pd2529-LTBP-10×His (LTBP-1 ETBE or LTBP-3 ETBE) plasmids were co-transfected at a total DNA concentration of 2 μg/mL. For TβRII-Fc expression, pd2529-TβRII-Fc plasmid was transfected alone. Glucose was supplemented to a final concentration of 3 g/L at the time of transfection to enhance cell viability and protein yield. The conditioned media were harvested after 5 days’ culture by centrifugation at 4000 × g for 30 min.

For protein purification of the LTBP/pro-TGF-β complex, the conditioned media were first filtered through a 0.22 μm PES membrane. The clarified media were then directly mixed with Ni-NTA beads (Smart-Lifesciences) at 4 °C for 3 h. The beads were washed sequentially with Buffer A (20 mM Tris pH 8.0, 500 mM NaCl) and Buffer B (20 mM Tris pH 8.0, 500 mM NaCl, 25 mM imidazole) for 20 column volumes (CVs) each. The complex was eluted with Buffer C (20 mM Tris pH 8.0, 500 mM NaCl, 300 mM imidazole), followed by cleavage of the 10×His-tag via incubation with PreScission protease at 4 °C overnight. The eluate was then buffer-exchanged into Buffer D (20 mM Tris pH 8.0, 50 mM NaCl) and loaded onto a HiTrap Q HP anion exchange column (Cytiva) pre-equilibrated with the same buffer. The protein was eluted with a linear gradient of 50–500 mM NaCl over 20 CVs. Fractions containing the target complex were pooled and concentrated for the final polishing step by size exclusion chromatography (SEC) using a Superose 6 Increase 10/300 GL column (Cytiva) equilibrated with Buffer E (20 mM HEPES pH 7.5, 150 mM NaCl).

For TβRII-Fc purification, the filtered media were loaded onto a Protein A Sepharose FF column (Cytiva) pre-equilibrated with Buffer A (20 mM Tris pH 8.0, 500 mM NaCl). After washing with 20 CVs of Buffer A, the protein was eluted with 0.1 M glycine-HCl (pH 3.0) and immediately neutralized with 1 M Tris-HCl (pH 9.0). The eluate was then buffer-exchanged into Buffer E and subjected to SEC using the same Superose 6 Increase 10/300 GL column equilibrated with Buffer E.

### Cryo-EM sample preparation and data collection

The purified LTBP-1/pro-TGF-β1 and LTBP-3/pro-TGF-β3 complexes were concentrated to approximately 5 mg/ml for cryo-EM grid preparation. Aliquots of 3.5 μl protein sample were applied to glow-discharged holey carbon grids (Quantifoil R1.2/1.3 Au 300). The grids were blotted for 3 s under 100% humidity at 8 °C using a Vitrobot Mark IV (FEI) and subsequently vitrified by plunge-freezing into liquid ethane cooled by liquid nitrogen.

Cryo-EM data were collected on a 300-kV Titan Krios microscope (Thermo Fisher) equipped with a K3 direct electron detector (Gatan) and controlled by EPU software. Micrographs were recorded at a nominal magnification of 81,000 ×, corresponding to a physical pixel size of 1.07 Å per pixel, with a nominal defocus range of 1.0-2.0 μm. Each movie comprised 40 frames with a total exposure time of 3.2 s, yielding a total electron exposure dose of approximately 58.7 e⁻/Å². The data collection parameters are summarized in Supplementary Table 1.

### Cryo-EM data processing

The image stacks were gain-normalized and corrected for beam-induced motion using MotionCor2^53^ in RELION 4.0. Contrast transfer function (CTF) parameters^54^ were estimated using CTF Estimation. Micrographs exhibiting ice contamination or CTF fitting worse than 4 Å were discarded. High-quality micrographs were subjected to auto-picking, and the extracted particles underwent multiple rounds of 2D classification to remove junk particles, aggregates, and incomplete complexes. Particles belonging to well-defined structural classes were retained and subsequently imported into cryoSPARC^55^ (version 3.2.0) for 3D reconstruction and refinement.

For the LTBP-1/pro-TGF-β1 complex, approximately 992,100 particles were selected for ab initio reconstruction and heterogeneous refinement. Particles from well-defined classes were combined, and duplicate particles were removed. Further iterative rounds of 3D classification with subsequent ab initio reconstructions and heterogeneous refinements were conducted to eliminate suboptimal particles. The final selected particle set comprising 160,694 particles was subjected to non-uniform (NU) refinement, yielding a 3.6 Å resolution map determined by the gold-standard Fourier shell correlation (FSC) = 0.143 criterion.

For the LTBP-3/pro-TGF-β3 complex, a similar processing workflow was employed starting from approximately 1.9 million initial particle images. Following iterative ab initio reconstruction, heterogeneous refinement, and duplicate removal, 523,744 final particles were selected for NU refinement, resulting in a 3.4-Å resolution map at the FSC = 0.143 threshold. Detailed data processing workflows are presented in Supplementary Figures 1 and 2.

### Model building and refinement

The initial structural models for LTBP-1/pro-TGF-β1 and LTBP-3/pro-TGF-β3 were generated by docking available high-resolution crystal structures and AlphaFold-predicted models into the cryo-EM density maps using UCSF ChimeraX^56^. For pro-TGF-β1, the previously reported structure of the LRRC33/pro-TGF-β1 complex^10^ (PDB: 7Y1R) and the unbound LTBP-1 TB domain structure ^57^(PDB: 1KSQ) were used as initial templates. For pro-TGF-β3, the pro-TGF-β3/GARP complex structure ^58^ (PDB: 8VSB) and the LTBP-3 TB domain AlphaFold model (AF-O35282-F1-v6)^59^ were employed as starting models. The fitted models were subsequently subjected to iterative rounds of real-space refinement in PHENIX^60^ with secondary-structure and geometric constraints, interspersed with manual inspection and adjustment of side-chain rotamers and backbone conformations in Coot^61^. Alternative automatic and manual refinements were performed until convergence, with particular attention paid to the disulfide bond connectivity at the covalent interface and the loop-flipping-mediated hydrophobic interaction regions. The geometries of the final structural models were validated using MolProbity, and the statistics of Ramachandran plots were examined. The final models were validated against the respective maps by model-to-map FSC calculations.

### Surface plasmon resonance (SPR) analysis

SPR experiments were performed using a Biacore T200 system (Cytiva) at 25 °C. A general amine coupling protocol was employed to immobilize various ligand proteins, including full-length or truncated LTBP variants and their corresponding mutants, TβRII-Fc and TβRI-Fc onto CM5 chips, aiming for an immobilization level of 200 resonance units (RU). A reference flow cell was left blank to allow double-referencing subtraction. Purified analytes, including pro-TGF-β isoforms (β1, β2, β3), their specified mutants, and various LTBP truncation fragments, were prepared in a series of indicated concentrations (0.1-10 μM) and injected sequentially at a flow rate of 20 μl/min in HBS buffer (20 mM HEPES pH 7.5, 150 mM NaCl, and 0.05% Tween-20). The sensor surface was regenerated using 10 mM glycine-HCl pH 1.5 by a 30-s pulse (50 μl/min) at the end of each cycle to restore the resonance units to the baseline. Sensorgrams were double-referenced and kinetic analyses were performed using Biacore T200 Evaluation Software. The experimental data were fitted to a 1:1 Langmuir binding model to generate the kinetic parameters. Fitting quality was assessed by χ² and R_max_ values.

### Biolayer Interferometry (BLI) analysis

Real-time, non-covalent binding kinetics were measured by biolayer interferometry (BLI) using an Gator® Prime system (Gator Bio) at 25 °C with continuous shaking at 1000 rpm. All binding assays were performed in a kinetic buffer consisting of PBS, pH 7.4, supplemented with 0.1% (w/v) BSA and 0.02% (v/v) Tween-20] to minimize non-specific interactions.

To determine the binding specificities of LTBP-2 fragments, biotinylated LTBP-2 ETBE, EGF-like, or TBE domains were immobilized onto pre-equilibrated Gator® Streptavidin XT (SA-XT) biosensors (Gator Bio, Cat 20-5120) to a threshold capture level of approximately 1.0 nm. After a 60 s baseline equilibration step in the kinetic buffer, the loaded biosensors were immersed into wells containing serially diluted analytes (purified pro-TGF-β1, -β2, and -β3 at indicated concentrations ranging from 250 nM to 4 μM) for an association phase of 120-200 s. The biosensors were subsequently transferred into analyte-free buffer to monitor the dissociation phase for 120-200 s. Binding assays for pro–TGF-β1, -β2, -β3 and LTBP-3/ pro–TGF-β3 to type I receptor (TβRI) were performed by immobilizing TβRI on the SA–XT sensor chip via biotin–streptavidin capture. Notably, the buffer consisted of analyte–free buffer containing 6 µM type I receptor (TβRII).

To correct for baseline drift and non-specific binding, all sensorgrams were double-referenced by subtracting the responses from a reference sensor (unloaded, dipped into analyte) and a reference well (ligand-loaded, dipped into buffer only). The association and dissociation responses were globally fitted to a 1:1 Langmuir binding model using Gator Data Analysis Software v2.0.

### Cell surface display of GPI-anchored LTBP/pro-TGF-β complexes and surface presentation analysis

Expi293F cells were cultured in FreeStyle 293 medium (Gibco) adherently at 37℃ with 5% CO_2_. Genes encoding LTBPs (full-length LTBP-1, -2, -3, -4, and specified domain truncations/mutants) were cloned into the pD2529 mammalian expression vector (ATUM). To anchor LTBP variants to the cell surface, a V5-tag and a GPI anchor sequence were fused to their C-termini. Genes encoding pro-TGF-β1, -β2, and -β3 (WT and mutants) were cloned into the pcDNA3.4 mammalian expression vector (Invitrogen). The “RRGDLAT” of pro-TGF-β1 and “GRGDLGRL” of pro-TGF-β3 were replaced with an HA-tag to prevent dissociation of mature TGF-β and to avoid interfering with complex formation (the N-terminus is involved in complex formation, thus N-terminal tagging is not optimal). Mutants were generated by two-step PCR using PrimeSTAR® HS DNA Polymerase (Takara, R044A) and cloned into Top10 Escherichia coli competent cells (Tsingke, TSC-C12) using the ClonExpress MultiS One Step Cloning Kit (Vazyme, C113-02). All mutations were validated by DNA sequencing.Expi293F cells transiently co-transfected with the indicated LTBP and pro-TGF-β plasmids were stained with an iFluor 488-conjugated anti-HA antibody (1 μg/ml, GenScript, Cat A01806, Clone 5E11D8) and an iFluor 647-conjugated anti-V5 antibody (1 μg/ml, GenScript, Cat A01805, Clone 11D5), and subjected to flow cytometry using a CytoFLEX cytometer (Beckman Coulter). The results were analyzed using FlowJo V10. Supernatants and total lysates of the transfected cells were subjected to non-reducing and reducing SDS-PAGE for immunoblotting with an anti-HA primary antibody (1:2000, Biolegend, Cat 901501, Clone 16B12) and an anti-Mouse IgG (whole molecule) HRP secondary antibody (1:30000, Sigma, Cat A9044) to detect pro-TGF-β and its specific complexes with LTBP variants.

### Pro-TGF-β activation in the ECM

To evaluate TGF-β signaling activation, 20,000 Expi293F cells transiently transfected with a TGF-β-SMAD3 responsive (CAGA)_12_-Luciferase reporter construct were seeded per well in 96-well plates. For soluble activation assays, the reporter cells were incubated with purified proteins across a range of concentrations to evaluate isofom-specific signaling. For integrin- and extracellular matrix (ECM)-dependent activation assays, cells were transfected with plasmids encoding LTBP variants and/or pro-TGF-β isoforms either in combination or individually (to serve as single-transfection controls). Notably, transient transfection of LTBP–2 and its 16th EGF–like domain deletion mutant consistently resulted in low protein expression, as determined by ELISA, which showed reduced concentrations in the extracellular matrix, precluding reliable activation measurements and comparison to the other isoforms. We therefore established stable Expi293F cell lines constitutively expressing full-length LTBP-2 or LTBP-2 EGF-like deletion mutant, and adjusted seeding densities to achieve expression levels comparable to those obtained by transient transfection of other LTBP family members. At 48 h post-transfection, the transfected cells were incubated with 20 mM EDTA for 1 min. The plates were then gently washed twice with PBS to completely detach and remove the cells while preserving the deposited ECM and its associated proteins on the plate. The relative abundance of anchored LTBP or LTBP/ pro-TGF-β complex in the ECM were measured by ELISA to ensure similar activation conditions. Subsequently, 5000 Expi293F cells transiently expressing specific integrins (α_v_β_6_- or α_v_β_8_-) were seeded together with 20,000 reporter cells onto this pre-deposited ECM.

After 24 h of incubation at 37 °C, cells were lysed for 30 min on ice using 50 μl per well 1× Passive Lysis Buffer (Promega, E1941). Then the supernatants were transferred into a 96-well solid white flat microplate (Corning, 3917). 100 μl luciferase substrate was added per well according to the manufacturer’s recommendations (Promega, E1501) and the chemiluminescence was measured using a Synergy H1 microplate Reader (BioTek).

### Enzyme-linked immunosorbent assay (ELISA)

Using the cell-derived ECM prepared as described in the pro-TGF-β activation assay, an in situ enzyme-linked immunosorbent assay (ELISA) was performed to quantify the relative abundance of anchored LTBP or variants and pro-TGF-β1.The ECM-coated wells were directly blocked with a blocking buffer containing 3% (w/v) non-fat dry milk and 2% (w/v) BSA in PBST (PBS containing 0.05% Tween-20) overnight at 4 °C. Following three washes with PBST, the wells were incubated with primary antibodies diluted in the blocking buffer (1:2000) for 1 h at 37 °C. Specifically, an anti-HA antibody (Biolegend, Cat 901501, Clone 16B12) was used to detect HA-tagged LTBP-2 and its variants, while an anti-TGF-β1 antibody (Clone 28G11) was used to detect pro-TGF-β1.

After three washes with PBST, the wells were incubated with an HRP-conjugated goat anti-mouse IgG secondary antibody (1:20,000, Sigma, Cat A9044) for 1 h at 37 °C. The plates were then rigorously washed five times with PBST. For colorimetric development, 100 μL of 3,3’,5,5’-tetramethylbenzidine (TMB) substrate solution was added to each well and incubated for 10 min at 37 °C in the dark. The reaction was terminated by the addition of 50 μL of 2 M H_2_SO_4_ stop solution. The optical density at 450 nm (OD450) was immediately measured using a Synergy H1 microplate reader (BioTek). Background signals obtained from mock-transfected ECM wells were subtracted for data analysis.

### Pull-down

His-tagged LTBP-2 truncation constructs were expressed in Expi293F cells either alone or by co-transfection with pro-TGF-β1 at a 1:1 plasmid mass ratio. Cells were transfected at a density of 3×10^6^ cells/mL and cultured for 5 days, after which the conditioned media containing the secreted proteins were collected as described above. His-tagged proteins and their co-associated binding partners were captured from the clarified supernatants by Ni-NTA affinity pulldown.

Ni-NTA pull-down eluates were resolved by SDS-PAGE under reducing (R) or non-reducing (NR) conditions and transferred to PVDF membranes. Membranes were blocked with 5% non-fat milk in TBST for 1 h at room temperature, incubated with primary antibody overnight at 4 °C, and then with HRP-conjugated secondary antibody for 1 h at room temperature. Non-reduced blots were probed with anti-pro-TGF-β1(clone 28G11, 1:2000) and reduced blots with anti-TGF-β1 (Abclonal, A2124, 1:2000); FLAG-tagged LTBP-2 proteins were detected with anti-FLAG antibody (1:2000). Goat anti-mouse IgG (whole molecule)-HRP (Sigma, A9044, 1:20000) ;and goat anti-rabbit IgG–HRP (Invitrogen, 31460,1:20000) were used as secondary antibodies. Signals were visualized by enhanced chemiluminescence. Co-precipitation of pro-TGF-β1 with a given LTBP-2 truncation in the Ni-NTA pull-down fraction indicates an interaction between the two proteins.

## Data availability

The cryo-EM density maps of LTBP1/pro-TGF-β1 complex and LTBP3/pro-TGF-β3 complex have been deposited in the Electron Microscopy Data Bank under the accession codes EMD-XXXXand EMD-XXXX.

## Acknowledgements

This research was funded by the National Natural Science Foundation of China (No. 32471261 to B.Z.; No. 32401010 to L.W. and No. 32471252 to Z.Z.), Shenzhen Science and Technology Program (No. JCYJ20220818102018038 to B.Z. and No. JCYJ20230807120215031 to F.T.), the Beijing Natural Science Foundation (No. Z240013 to Z.Z.), the Basic and Applied Basic Research Fundation of Guangdong Province (No. 2024A1515220026 to F.T.), the Fundamental Research Funds for the Central Universities (No. BMU2026YJ010 and No. BMU2026RCZX095 to Z.Z.) and the Young Project of Natural Science Exploration of Zhejiang Chinese Medical University (No. 2025JKZKTS06 to Y.L.). The study was also supported in part by the Center for Life Sciences and the Qidong-SLS Innovation Fund (to Z.Z.). We thank the Cryo-EM platform at the School of Life Sciences (SLS) of Peking University for help with cryo-EM data collection.

## Author contributions

Conceptualization, B.Z., Z.Z. Y.L. and F.T.; Methodology, D.Z., L.W. and X.L.; Investigation, L.W., D.Z., J.T., X.L., X.Z., P.Y., J.L., R.L., N.W. and Z.W.; Formal Analysis, L.W. and D.Z.; Writing Original Draft, B.Z. L.W. and J.T.; Writing – Review & Editing, B.Z. Z.Z.; Visualization, L.W., D.Z. and J.T.; Supervision, B.Z., Z.Z., Y.L. and F.T.; Funding Acquisition, B.Z., L.W., Z.Z., Y.L. and F.T.

## Declaration of Generative AI and AI-assisted technologies in the writing process

During the preparation of this work, the authors used a generative AI tool (DeepSeek) to assist with language polishing and grammatical refinement of the manuscript text. After using this tool, the authors carefully reviewed and edited the content as needed and take full responsibility for the final content of the publication.

